# The *Ex Vivo* Infection of the Peripheral Bovine Mononuclear Cells (PBMCs) and the Bovine Spleen Cells with the Bovine Coronavirus (BCoV) Induced a Differential Expression of the Host Cytokine Genes Profiles and Modulates the Virus Replication

**DOI:** 10.1101/2024.07.01.601600

**Authors:** Abid Ullah Shah, Maged H Hemida

## Abstract

The adaptive immune response during BCoV infection of peripheral blood mononuclear cells (PBMCs), the bovine spleen cells, and their isolated T lymphocytes was not studied well. Our study confirmed successful BCoV infection in PBMCs and spleen T cells. This was evidenced by measuring genome copy numbers using real-time PCR, expression levels of BCoV spike and nucleocapsid proteins via western blot and immunofluorescence assays, and virus infectivity titration by plaque assay. In infected PBMCs, CD4 T-cell levels were 1.45-fold higher, and CD8 T-cell levels were 1.6-fold lower compared to sham-infected cells. Conversely, infected splenocytes showed a 0.88-fold decrease in CD4 T-cells and a 1.88-fold increase in CD8 T-cells. The cytokine gene expression analysis revealed that BCoV infection activated type 1 interferon and upregulated IL-6 expression in PBMCs and splenocytes. These findings demonstrate that BCoV successfully infects immune cells from PBMCs and spleen, inducing differential host cytokine gene expression favors virus replication.

## 1 Introduction

Bovine coronavirus (BCoV) is one of the most prevalent viral threats for the cattle industry in both beef and dairy sectors (1, 2, 3). The BCoV infection is ubiquitous in young calves (up to 6 months of age) across the world (4). The BCoV is responsible for several clinical syndromes in the affected cattle, including diarrhea in young calves, winter dysentery, and respiratory manifestations of all ages, and it also participated in the development of the bovine respiratory diseases complex along with other viral, bacterial, and parasitic pathogens (4).

Although the roles of the peripheral blood mononuclear cells (PBMCs) and the bovine spleen cells in the innate and adaptive immune responses have been studied well in many viral diseases of cattle (5, 6, 7, 8), little is still known about the roles of these immune cells in the immune regulation of the BCoV infection. There is a scarcity of commercially available BCoV vaccines. Thus, studying the immune response against BCoV infection using various models (*in vitro*, *ex vivo,* and *in vivo*) may help develop some novel vaccines and antiviral therapies that help prevent and control BCoV infection in animals, respectively. The bovine peripheral blood mononuclear cells (PBMCs) play important roles in the immune response against most pathogens affecting cattle (9). The PBMCs consist of several types of cells, including B cells, T cells, N.K. cells, and monocytes. The cell-mediated immune response plays an important role during the BCoV infection in calves (10). There are several subpopulations of T cells. Each population produces some cytokines that shape the immune response against viral pathogens in cattle. The CD4 has four main subpopulations (Th1, TH2, TH17, and the Treg cells) (10, 11, 12, 13). The Th1 mainly produces IFN-γ and IL-2, which play important roles in the clearance of viral infections (14). The Th2 produces several cytokines, including IL-4, IL-5, IL-10, and IL-13, which stimulate B cell proliferation, which is mainly important in the immune response to extracellular pathogens and inflammatory reactions (15). The Th17 mainly produces both IL-17 and IL-22, which function mainly in response to bacterial and fungal infections (16, 17, 18). The Treg T cell mainly produces IL-10 and TGF-β anti-inflammatory cytokines (6, 19).

On the other hand, CD8 T cells are called cytotoxic T cells and mainly help destroy the viral infected cells to curb the viral infection in the affected host (20, 21, 22, 23, 24). The PBMCs are involved in the process of phagocytosis, the production of reactive oxygen, and the production of common cytokines and chemokines during viral infection in cattle. The PBMCs also tune the balance between the inflammation and tissue damage during viral infection in cattle, such as the bovine viral diarrhea virus (BVDV), the foot and mouth diseases virus (FMDV), bovine respiratory syncytial virus (BRSV), Bovine Herpesvirus-1 (BHV-1), (25, 26, 27, 28). The spleen is one of the major immune organs in cattle, and it plays an important role in the regulation of the immune response in cattle against major pathogens (9). The spleen is a multitasking organ during viral infection in any host, including cattle. It coordinates between the innate and adaptive immune response.

The spleen starts by filtration of the virus from the blood and tries to eliminate the virus from the bloodstream. The macrophages and dendritic cells in the spleen reduce the number of viral particles by engulfing the viral infected cells. This is in addition to their roles as antigen-presenting cells of the viral proteins to the major histocompatibility complex and subsequently activating T cells. The spleen CD4 cells produce several cytokines to stimulate the B cells and CTL action. The spleen CD8 is also differentiated into CTL to eliminate the viral infected cells (7, 29, 30).

The CD4/CD8 ratio has several implications for the prognosis, diagnosis, and monitoring of the therapeutic progression of many viral infections in humans and animals (31, 32, 33, 34, 35, 36). The balanced CD4/CD8 ratio indicates that the immune system is normal. If the CD4/CD8 ratio is high (the CD4 population is much higher than the CD8 population). In that case, this pattern suggests the predomination of the T helper cell, which potentiates the humoral immune response and antibody production. If the CD4/CD8 ratio is small, that means (the CD8 population is higher than the CD4); this indicates a robust cytotoxic immune response. This phenomenon is widespread at the end of the course of the viral infection, in which the host tries to clear the viral infected cells. A significant reduction in the CD4/CD8 ratio during BVDV infection suggests immunosuppression (37, 38).

Although the roles of different immune cells have been studied in many viral diseases of cattle, particularly the BVDV, BHV-1, FMDV, BRSV, and BLV. (9, 19, 24, 25, 26, 28), the BCoV immune response, especially cell-mediated immunity, has not been studied well. The main goals of the current study were to study the roles of different immune cells in the bovine PBMCs and spleen in the response to BCoV infection using the *ex vivo* models, particularly the T lymphocytes isolated from the PBMCs and spleen infected with either the enteric or respiratory isolates of BCoV.

## 2. Materials and Methods

### 2.1. Viruses

The Bovine Coronavirus (BCoV) enteric isolate (Accession Number: U00735) was obtained from B.E.I. Resources (NIAID, N.I.H.; BCoV, Mebus, NR-445). The BCoV respiratory isolate (Accession Number: ON146444) was generously provided by Dr. Aspen Workman from the Animal Health Genomics Research Unit at the USDA Agricultural Research Service, U.S. Meat Animal Research Center (39).

### 2.2. Isolation of the PBMC and BCoV infection protocol

The peripheral blood mononuclear cells (PBMC) were isolated from bovine whole blood obtained from Lampire Biological Laboratories (New York, U.S.A.). The blood was collected from healthy donors and mixed with 2% sodium EDTA. Blood was tested for the presence of BVDV and mycoplasma. PBMC were isolated by gradient centrifugation using Histopaque-1077 (Sigma; Lot. No. RNBL7068) following the manufacturer’s instructions. The PBMCs were lysed with Red Blood Cell (R.B.C.) Lysing Buffer Hybri-Max (Sigma; Cat. No. R7757-100ML) to remove the R.B.C.s completely from the PBMCs. After washing, PBMCs were cultured in RPMI-1640 (ATCC, 30-2001) supplemented with 10% horse serum (Gibco; Ref. No. 26050-088) and 1% 10,000 ug/mL streptomycin and 10,000 units/mL penicillin antibiotics (Gibco; Ref. No. 15140-122) at 37°C in 5% CO_2_ for 24 hours. The next day, PBMCs were infected with 1 M.O.I. of BCoV/Ent or BCoV/Resp isolate for two hours and incubated at 37°C in 5% CO_2_. The PBMCs were cultured for 72 hours at 37°C in 5% CO_2_ and then subjected to subsequent experiments.

### 2.3. Isolation of the bovine splenocytes and BCoV infection protocol

Spleen samples were collected from fresh, slaughtered young and healthy calves. The spleen samples from each bovine were tested for BCoV infection. Single-cell splenocyte suspension from spleen samples was prepared as described previously (40). Briefly, the spleen was minced using sterile scissors and filtered through a 70-um nylon mesh sterile cell strainer (Fisherbrand; Cat. No. 22363548). The resultant splenocytes were suspended and washed three times with sterile PBS containing 2% streptomycin/penicillin antibiotics. The pellet was incubated at room temperature with the RBCs lysis buffer according to the manufacturer’s instructions to remove the red blood cells and centrifuged at 400 x g for 5 minutes at 4°C. The pellet was resuspended in RPMI-1640 containing 10% horse serum and 1% streptomycin/penicillin antibiotics filtered through a 70-um sterile cell strainer and incubated at 37°C in 5% CO_2_. After 24 hours, splenocytes were infected with (MOI=1) of BCoV/Ent or BCoV/Resp isolate for two hours and incubated at 37°C in 5% CO_2_. The splenocytes were cultured for 72 hours at 37°C in 5% CO_2_ and then subjected to subsequent experiments.

### 2.4. Isolation and cultivation of the T-lymphocytes from PBMCs and Splenocytes

The purified PBMCs were cultured in RPMI-1640 containing 10% horse serum, 1% streptomycin/penicillin antibiotics, and 1 ug/mL phytohemagglutinin (PHA.) (Gibco; Ref. No. 10576-015) for 24 hours at 37°C in 5% CO_2_. This step will allow monocytes to adhere to the culture plate, and lymphocytes in the PBMCs remain in the suspension. The next day, the supernatant was removed and centrifuged at 500 x g for 5 minutes. The pellet containing lymphocytes was resuspended in RPMI-1640 containing 10% horse serum, 1% streptomycin/penicillin antibiotics, and 1 ug/mL PHA. After incubation for 72 hours at 37°C in 5% CO_2_, the supernatant was centrifuged at 500 x g for 5 minutes. The pellet was resuspended in RPMI-1640 containing 10% horse serum, 1% streptomycin/penicillin antibiotics, and 20 ng/mL of Recombinant Bovine IL-2 (KingFisher Biotech, Inc; Lot. No. HS3386ES) at 37°C in 5% CO_2_. The recombinant bovine IL2 will allow the T lymphocytes to grow in the media. After 1-2 days, the supernatant was collected and centrifuged at 500 x g for 5 minutes, and the pellet was infected with 1 M.O.I. of BCoV/Ent or BCoV/Resp isolate for two hours. The cells were gently homogenized every 30 minutes during infection. Cells were washed and incubated at 37°C in 5% CO_2_ for 72 hours. The supernatant was centrifuged at 500 x g for 5 minutes, and the pellet containing T lymphocytes was resuspended in media containing 1mg/mL of D-glucose and subjected to subsequent experiments.

### 2.5. The RNA extraction and quantitative real-time-PCR (qRT-PCR)

The total RNAs were isolated from control (sham), BCoV/Ent, and BCoV/Resp-infected PBMCs, splenocytes, and T lymphocytes using TRIzol LS Reagent (Invitrogen; REF: 10296010) following the manufacturer’s instructions. The RNA concentrations and 260/280 ratios were assessed using a NanoDrop OneC (Thermo Scientific). The extracted RNAs were then converted into cDNA using a high-capacity reverse transcription kit (Applied Biosystems; Lot: 2902953), according to the manufacturer’s protocol. Quantitative reverse transcription polymerase chain reaction (qRT-PCR) was performed using PowerUp SYBR Green Master Mix (Applied Biosystems; Lot: 2843446) as per the manufacturer’s instructions and analyzed with a QuantStudio3 (Applied Biosystems). The oligonucleotides used for qRT-PCR assays were designed using the Primer3 online tools (https://primer3.org/) (41). The gene expression levels and BCoV expression were normalized to the bovine β-actin using the 2^−ΔΔCt method (42). All oligonucleotides used in this study and the relevant information are listed in Table 1.

**Table 1:**
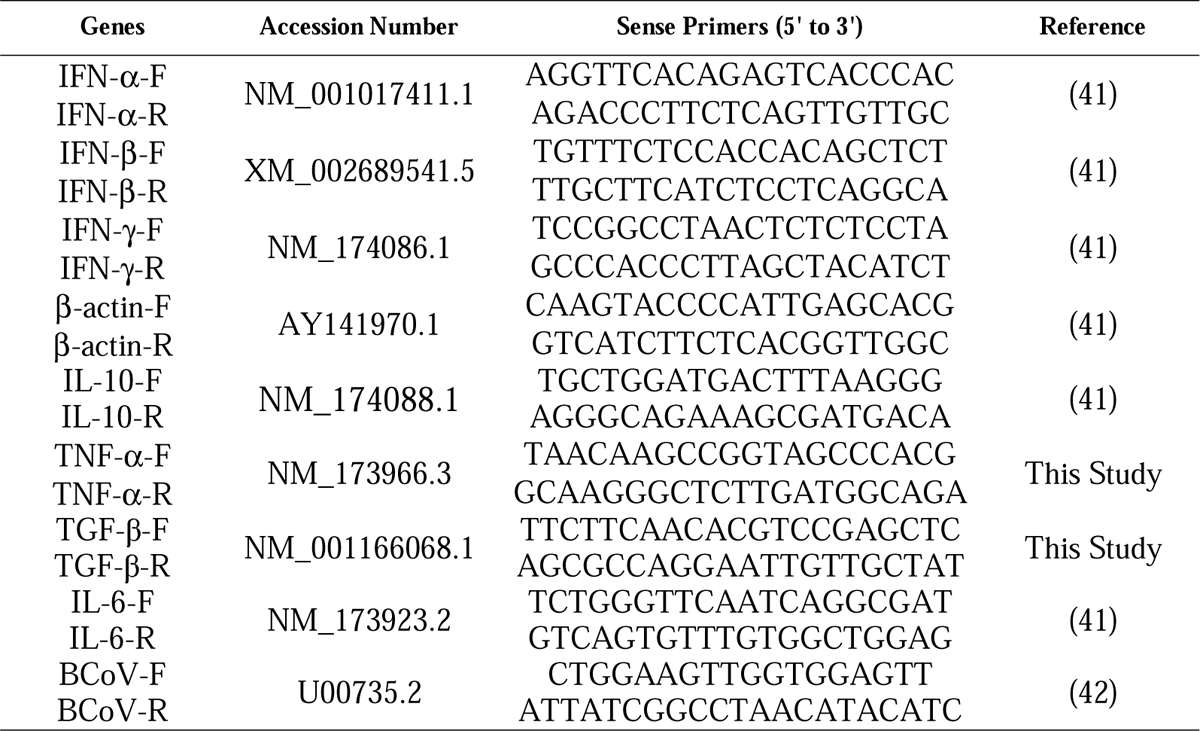
List of the oligonucleotides used for the amplification of the BCoV and host cytokine genes.

### 2.6. The Western Blot technique

Either the PBMCs or the splenocytes were centrifuged at 500 x g for 5 minutes, and the pellet was washed with ice-cold phosphate-buffered saline (PBS). The cells were then lysed with radioimmunoprecipitation assay (RIPA) lysis buffer (Thermo Scientific; REF: 89901), supplemented with 1% 0.5 M EDTA solution and 1% protease and phosphatase inhibitor (Thermo Scientific). The lysate was centrifuged at 14,000 rpm for 15 minutes, and the supernatant was used to measure protein concentration with a BCA. kit (Thermo Scientific) according to the manufacturer’s instructions. The protein concentrations were quantified using a NanoDrop OneC (Thermo Scientific). An equal volume of 2x Laemmli buffer 2x laemmli sample buffer (Bio-Rad; Cat. No. 1610737) was added to the lysate, and the mixture was incubated at 100°C for 10 minutes. The proteins were then separated by electrophoresis on a 10% SDS-polyacrylamide gel and transferred to a polyvinylidene difluoride (PVDF) membrane (Bio-Rad, Cat. No. 1620177). The membrane was blocked with 5% bovine serum albumin (B.S.A.) in Tris-buffered saline (TBS) containing 0.05% Tween-20 (TBST) and incubated overnight at (4°C) with primary antibodies specific to the proteins of interest. The following day, the membrane was washed with TBST and incubated with horseradish peroxidase (HRP)-conjugated secondary antibodies in the blocking reagent for 1 hour at room temperature. After additional washing with TBST, the membrane was soaked with Clarity Western Enhanced Chemiluminescence (ECL) Substrate (Bio-Rad; Cat. No. 170-5060), and Western blot bands were visualized using the GelDoc Go Imaging System (Bio-Rad). The primary antibodies used were: (BCoV-Nucleocapsid mouse anti-bovine monoclonal (Clone: FIPV3-70; Cat. No. MA1-82189), the BCoV-Spike rabbit anti-bovine polyclonal (Cat. No. PA5-117562), and the β-actin rabbit anti-bovine polyclonal (Cat. No. PA1-46296)), all these antibodies were purchased from Invitrogen. The secondary antibodies used in this study were the horseradish peroxidase (HRP)-conjugated IgG (H+L), the goat anti-rabbit (REF: 31460), and the goat anti-mouse (REF: 31430), also were purchased from Invitrogen.

### 2.7. The Immunofluorescence Assay

The PBMCs, splenocytes, and their isolated T lymphocytes were fixed in 80% acetone (Sigma; Cat. No. 179124)) for 10 minutes at (37°C) followed by centrifugation at 500 x g for 5 minutes. Next, the cells were incubated with ice-cold methanol for 10 minutes at (−20°C) followed by centrifugation at 500 x g for 5 minutes. The cells were permeabilized by resuspending cells with 0.5% Triton X-100 in 1 x PBS and incubated at room temperature for 15 minutes, followed by centrifugation at 500 x g for 5 minutes. Subsequently, cells were blocked with 2% B.S.A. in 1 x PBS at room temperature for 60 minutes. Without washing, the cells were incubated in primary antibody BCoV-Nucleocapsid mouse anti-bovine monoclonal (Clone: FIPV3-70; Cat. No. MA1-82189) purchased from Invitrogen diluted in 0.5% B.S.A. (1:100) overnight at 4°C in the dark. The next day, cells were washed with 1 x PBS and incubated with the goat anti-mouse (REF: 31430) secondary antibody conjugated with Alexa-Flour 488 obtained from Invitrogen desired concentration (1:200) for 1 hour at room temperature in the dark. For the co-localization and double staining, the cells were resuspended with the CD3 T-cell antibody conjugated with Brilliant Violet 605 (Clone: 17A2; Cat. No. 100237) purchased from BioLegend at the desirable concentration (1:100) diluted in 0.5% BSA for 2 hours at room temperature in the dark. The cells were counterstain with DAPI (Invitrogen; Ref. No. D1306) for 5 minutes at room temperature and examined immediately under a fluorescent microscope (ZEISS). Cells were washed with 1 x PBS after each step.

### 2.8. The Flow Cytometry assay and the data analysis

The PBMCs, splenocytes, and the T-lymphocytes from sham and the BCoV infected groups were equally divided (1 × 10^6^) into different groups. The cells were washed with PBS and resuspended in primary antibody of mouse anti-bovine CD4 (Clone IL11A; isotype IgG2a), mouse anti-bovine CD8 (Clone BQ1A; Isotype IgM), or mouse anti-bovine IFN-γ (Clone BQ1A; Isotype IgM) at desirable concentration (1ul/1 × 10^6^ cells) diluted in 100ul 1 x PBS for 1 hour at 4°C in the dark. All the primary antibodies were purchased from the Washington State University (WSU) Monoclonal Antibody Center (Washington, USA). Subsequently, cells were washed with 1 x PBS and incubated with the goat anti-mouse (REF: 31430) secondary antibody conjugated with Alexa-Flour 488 at the desired concentration (0.5ul/1 × 10^6^ cells) for 1 hour at 4°C in the dark. To evaluate the cells’ viability, each sample was counterstained with the desired concentration (0.5ul/1 × 10^6^ cells) of 7AAD viability staining solution (Biolegend; Cat. No. 420404) before analysis. Finally, cells were analyzed using a Fluorescence-Activated Cell Sorter (FACS) (CYTEK; NL-2000). Cells were analyzed using forward and side scatter to identify live cells. Doublet discrimination was applied to ensure single-cell analysis. At least 10,000 events were collected for each sample. Data were analyzed using FlowJo V10 software (FlowJo, China). All results were expressed as mean fluorescence intensity (MFI) or cell percentage.

### 2.9. Statistical analysis

All results are reported as mean values ± standard deviation (S.D.), with statistical analyses conducted using GraphPad Prism v9. For comparisons between different groups, a one-way analysis of variance (ANOVA) was performed, followed by Tukey’s or Dunnett’s post-hoc test as appropriate. Statistical significance was determined with P values less than 0.05 (P < 0.05). In the graphical representations, statistical significance is indicated as follows: * P < 0.05, ** P < 0.01, *** P < 0.001, and **** P < 0.0001. The presented data are combined from at least three independent experiments unless otherwise specified.

## 3. Results

### 3.1. The *Ex vivo* model revealed the Bovine coronavirus (BCoV) successfully infects and replicates in the Peripheral Blood Mononuclear Cells (PBMCs) and their isolated T lymphocytes

The bovine PBMCs were infected with BCoV Enteric (BCoV/Ent) and BCoV Respiratory (BCoV/Resp) isolates at a multiplicity of infection (MOI=1) for 72 hours. The real-time PCR demonstrated that BCoV/Ent and BCoV/Resp isolates significantly infect the bovine PBMCs as demonstrated by high genome copy numbers (Fig. 1A). The BCoV/Ent infected PBMCs showed higher genome copy numbers (2.16-fold) higher than the BCoV/Resp infected PBMCs (Fig 1A). BCoV replication was also confirmed on the protein level expression, including BCoV-S glycoprotein and the BCoV-N protein by Western blot (Fig 1B). The high expression level of the BCoV-S protein was prominent in the case of the BCoV/Ent isolate infected cells (Fig 1B). A high viral genome copy number in the T-T-cells isolated from the PBMCs and infected independently with either BCoV/Ent/BCoV/Resp isolates compared to the sham infected T cells, which showed no amplification (Fig 1C). The BCoV genome copy numbers in the case of the T cells infected with the BCoV-Ent isolates were 8.82 fold higher than those T-cells infected with the BCoV/Resp isolate (Fig 1C).

**Fig. 1.**
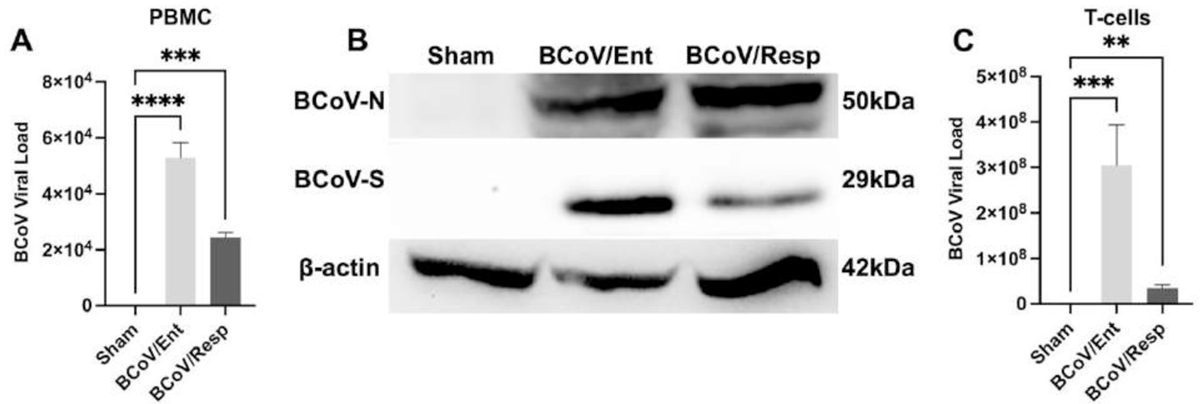
BCoV Infects the Bovine Peripheral Blood Mononuclear Cells (PBMCs) and their isolated Bovine T Lymphocytes. (A) Quantitative Reverse Transcriptase-Polymerase Chain Reaction (qRT-PCR) analysis of BCoV genomic viral load in PBMCs after 72 hpi; (B) Western blot analysis of BCoV-nucleocapsid (BCoV-N) protein, BCoV-spike (BCoV-S) protein, and β-actin protein expression in PBMCs after 72 hpi. (C) qRT-PCR analysis of BCoV viral load on T-cells isolated from bovine PBMCs.

### 3.2. Confirmation of the BCoV infection/replication in the PBMCs and their isolated T cells by the immunofluorescence

BCoV infection in the PBMCs was observed in the BCoV/Ent and BCoV/Resp infected groups compared to the sham-infected PBMCs by the immunofluorescence assay using the BCoV-N fluorescent conjugated antibodies (Fig. 2A – 2F). Our data shows a more prominent fluorescence signal in the case of the PBMCs infected with the BCoV/Ent isolate than those PBMCs infected with the BCoVC/Rep isolate (Fig. 2B, 2F). The doubled stained (merge) signals showed that a high level of BCoV/N protein was expressed in the case of the BCoV/Ent group, compared to BCoV/Resp infected PBMCs (Fig. 2C, 2F). The isolated T cells from the PBMCs showed a marked BCoV-N expression in both BCoV/Ent and BCoV/Rep isolates infected T cells (Fig. 2G – 2N). Our results also show that the red fluorescent signal associated with the CD3 T lymphocytes was prominently associated with the BCoV infection (Fig, 2H, 2L). The double staining showed that BCoV infects and co-localizes with most of the CD3 T lymphocytes present in the bovine PBMCs. The BCoV infection levels in the bovine CD3 T lymphocytes were similar in both BCoV/Ent and BCoV/Resp groups (Fig. 2J, 2N).

**Fig. 2.**
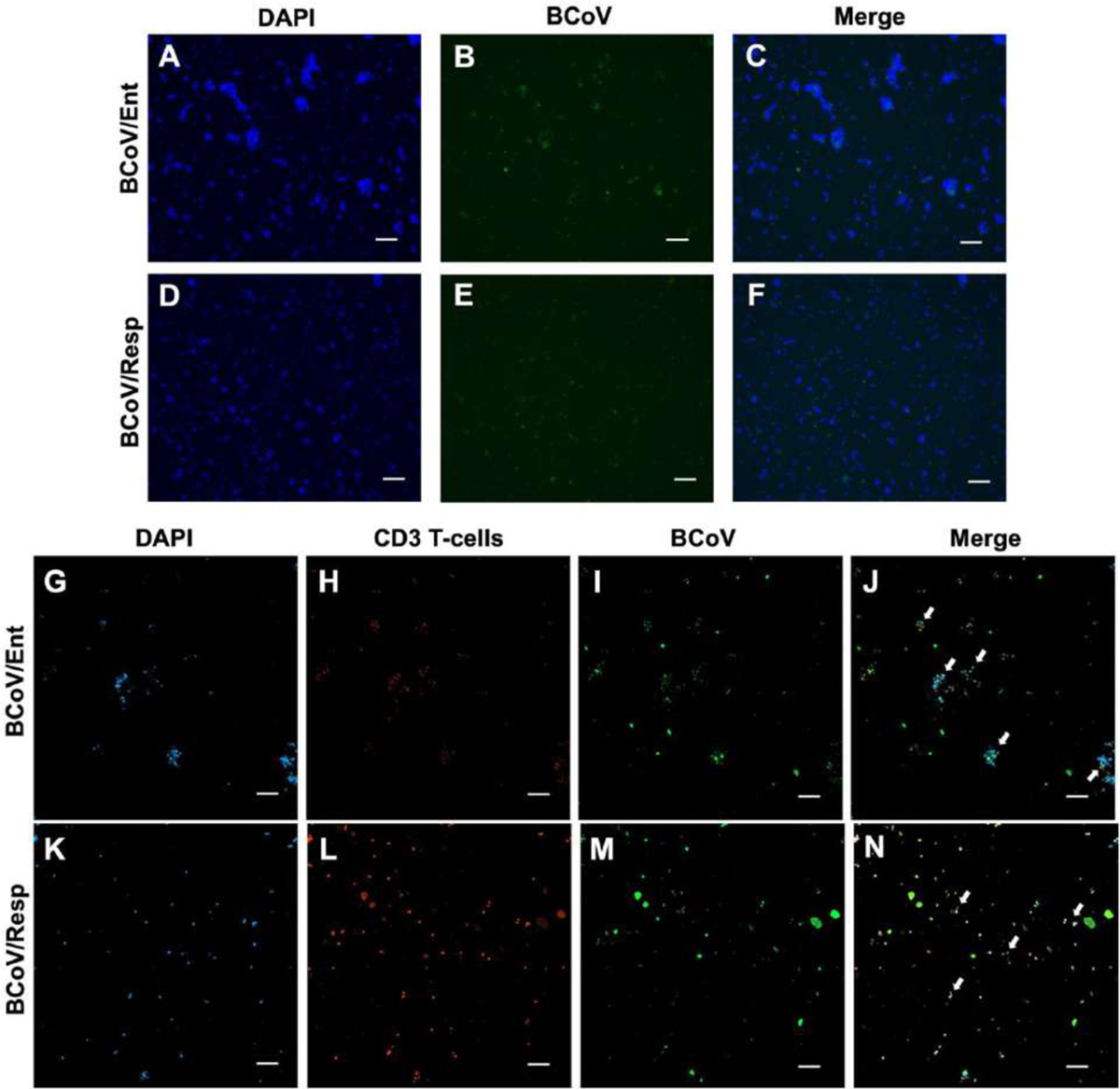
Immunofluorescence (IF) Analysis of the BCoV infection in the Bovine PBMCs and their isolated T-cells. (A-C) IF staining of PBMCs with (A) DAPI (blue color), (B) BCoV (green color), and (C) Merge in BCoV/Ent group. (D-F) IF staining of PBMCs with (D) DAPI, (E) BCoV, and (F) Merge in BCoV/Resp group. (G-J) IF detection of T-cells with BCoV/Ent isolate: (G) staining with DAPI, (H) CD3 T-cells (red color), (I) BCoV, (J) and the merged image showing co-localization (white arrowheads) of T-cells with BCoV. (K-N) IF detection of T-cells with BCoV/Resp isolate: (N) staining with DAPI, (L) CD3 T-cells (red color), (M) BCoV, (N) and the merged image showing co-localization (white arrowheads) of T-cells with BCoV. All the IFA images were captured at 10x magnification.

### 3.3. BCoV infection in PBMCS modulates the proliferation of T-cell sub-populations and the cytokines gene expression levels

To further explore the impacts of the BCoV infection on the subpopulation of T lymphocytes, the sham and the BCoV/Ent or the BCoV/Resp infected PBMC collected samples were analyzed using flow cytometry. The histogram representing the fluorescence intensity of the CD4 T-cells demonstrated a notable shift to the right in both BCoV/Ent and BCoV/Resp infected groups (Fig 3A). The Mean Fluorescence Intensity (MFI.) of the CD4 T-cells was significantly higher in both BCoV-infected groups compared to the sham group (Fig 3B). The CD8 T-cell population exhibited a distinct shift to the left in both infected groups (Fig. 3C). The M.F.I. of the CD8 T-cells was also significantly lower in both the BCoV/Ent and BCoV/Resp groups compared to the sham group (Fig. 3D), suggesting suppression of the CD8 T-cells in response to BCoV infection. The percentage of cells expressing CD4 and CD8 T-cells was inconsistent in sham and infected groups (Fig. 3G). The CD4 population was higher in the BCoV/Ent group (about 1.45 fold change), while the CD8 T-cells were 1.6 folds lower than that sham group (Fig. 3G). In the case of the BCoV/Resp group, both the CD4 and CD8 T-cell levels were downregulated compared to the sham group (Fig. 3G). To check the cytokine response of the BCoV infection in PBMCs, the IFN-γ expressions were analyzed. Results showed that the IFN-γ was significantly downregulated in the case of the BCoV/Ent infected group (Fig. 3E, 3F). These results indicate that BCoV/Ent infection was more severe in PBMCs, which activated host CD4 T helper cells, inhibited CD8 cytotoxic T-cells, and downregulated IFN-γ.

**Fig. 3.**
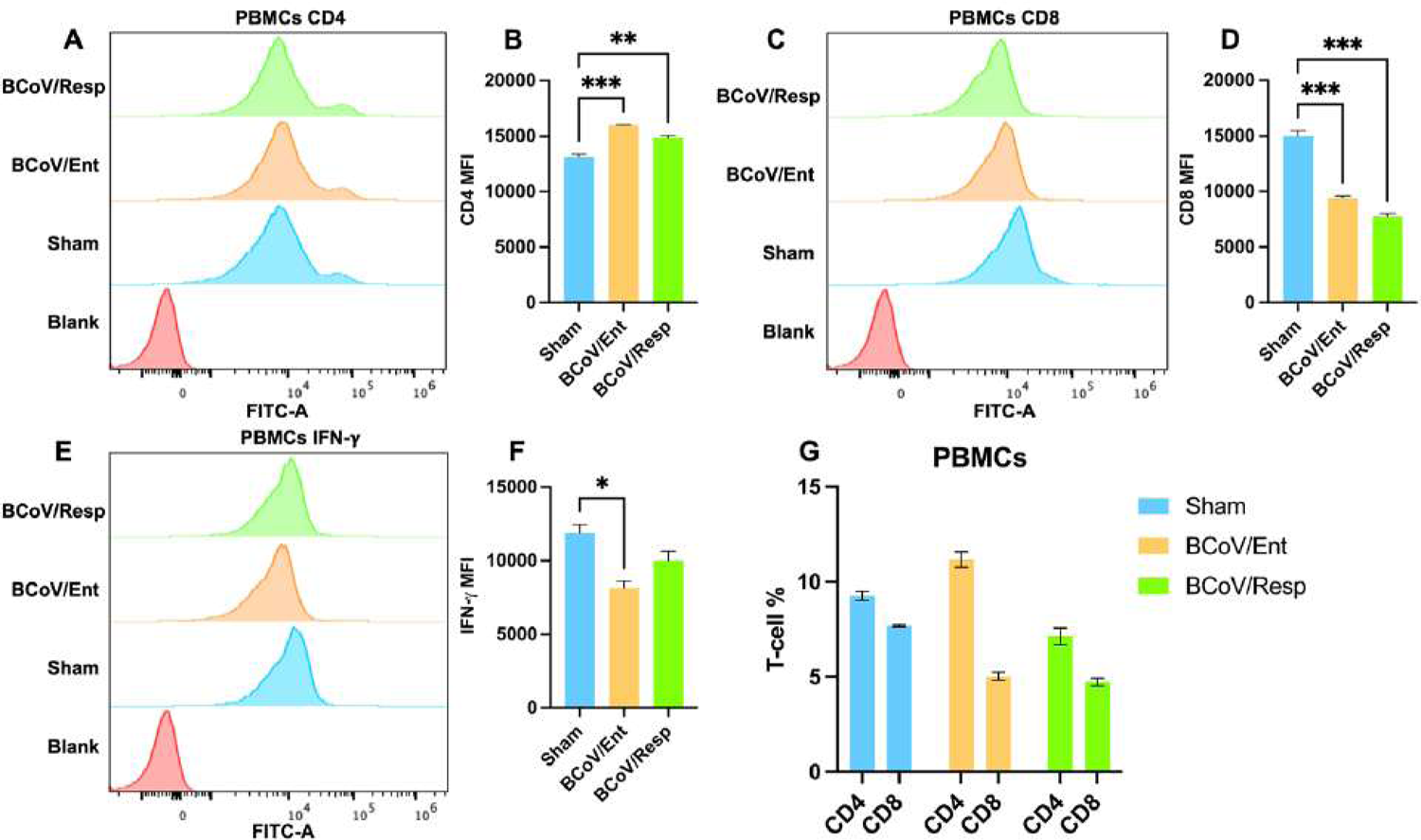
The impacts of BCoV/Ent and BCoV/Resp infection on the CD4, the CD8, and the IFN-**γ** expression levels in PBMCs. (A) Flow cytometry analysis showing histogram and (B) Mean fluorescence Intensity (MFI.) of CD4 T-cell population in sham, BCoV/Ent, and BCoV/Resp groups in PBMCs. (C) Flow cytometry analysis showing histogram and (D) MFI. of CD8 T-cell population in sham, BCoV/Ent, and BCoV/Resp groups in PBMCs. (E) Flow cytometry analysis showing histogram and (F) MFI. of IFN-γ in sham, BCoV/Ent, and BCoV/Resp groups in PBMCs. (G) CD4 to CD8 ration in sham, BCoV/Ent, and BCoV/Resp groups in bovine PBMCs.

### 3.4. BCoV Infection induced differential regulation of the CD4/CD8 T-lymphocytes Ratios using the *ex-vivo* model

To further validate the impacts of the BCoV infection on the T lymphocyte sub-population under the ex-vivo conditions, the total T lymphocytes were isolated from the bovine PBMCS infected with either the BCoV/Ent or the BCoV/Resp isolates compared to the sham infected group then and the obtained results were analyzed using the flow cytometry. Our results demonstrated a noticeable increase in the histogram peak of the CD4 T-cells in the BCoV/Ent group (Fig. 4A). The Mean Fluorescence Intensity (MFI.) of the CD4 T-cells was also significantly higher in the case of the BCoV/Ent infected group compared to the sham group (Fig 4B). The CD8 T-cell population exhibited a distinct shift to the right in both BCoV-infected groups (Fig. 4C). The M.F.I. of the CD8 T-cells was also significantly higher in both the BCoV/Ent and the BCoV/Resp-infected groups, compared to the sham infected group (Fig. 4D), suggesting the upregulation of CD8 T-cells in a response to BCoV infection. The percentage of cells expressing CD4 and CD8 T-cells ratio showed inconsistency following the BCoV infection (Fig. 4G). Remarkably, the CD4 T-cell percentage increased in the case of the BCoV/Ent infected group, while the CD8 T-cell percentage decreased compared to the sham-infected group (Fig. 4G). In the BCoV/Resp group, the CD4 and the CD8 T-cell ratio was consistent with the sham-infected group (Fig. 4G). The IFN-γ expression was significantly upregulated in the BCoV/Ent group, while no change was observed in the BCoV/Resp group, compared to the sham (Fig. 4E, 4F). Our results showed consistent CD4 and CD8 responses to the BCoV/Ent infection on PBMCs and their isolated purified T lymphocytes.

**Fig. 4.**
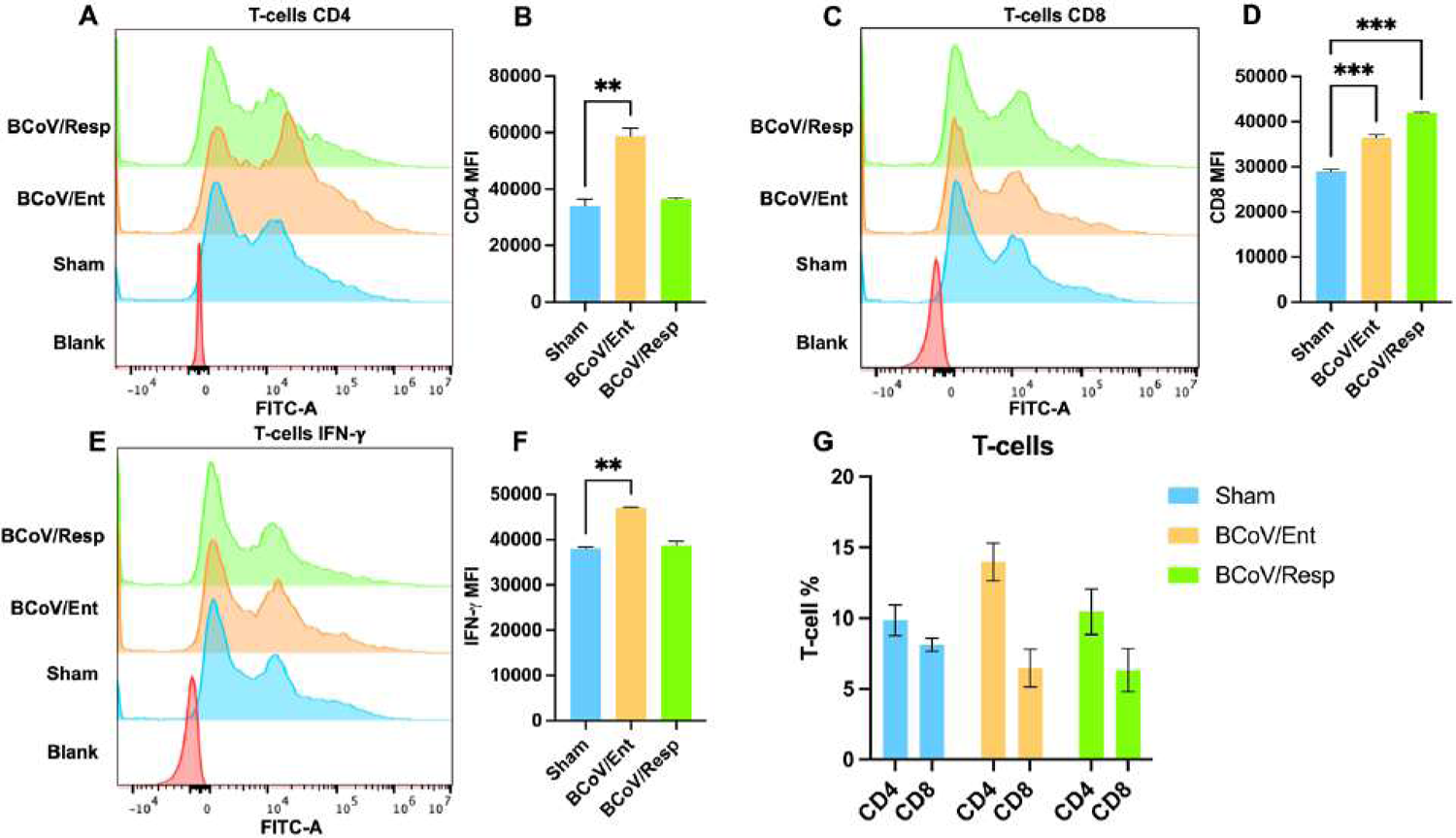
Evaluation of the CD4, the CD8, and the FN-**γ** production in the bovine T lymphocytes isolated from PBMCs after BCoV/Ent and the BCoV/Resp infection. (A) Flow cytometry histogram and (B) MFI. of CD4 T-cell in sham, BCoV/Ent, and BCoV/Resp groups in bovine T lymphocytes. (C) Flow cytometry histogram and (D) MFI. of CD8 T-cell in sham, BCoV/Ent, and BCoV/Resp groups in bovine T lymphocytes. (E) Flow cytometry histogram and (F) MFI. of IFN-γ in sham, BCoV/Ent and BCoV/Resp groups in bovine T lymphocytes. (G) CD4 to CD8 ration in sham, BCoV/Ent, and BCoV/Resp groups in bovine PBMCs.

### 3.5. The *ex vivo* model of the BCoV infection in the PBMCs revealed the activation of the cytokine-like storm

Results revealed a significant upregulation of the IFN-β and the IFN-γ genes expression in both the BCoV/Ent and the BCoV/Resp infected groups of the PBMCs, compared to sham, indicating a robust activation of antiviral responses and innate immune activation (Fig. 5B, 5C). The pro-inflammatory cytokines IL-6 and TNF-α expressions were significantly upregulated in both BCoV/Ent/Resp PBMCS infected groups, indicating the inflammatory nature of the host response during BCoV replication (Fig. 5D, 5F). The anti-inflammatory cytokine IL-10 also showed a significant upregulation in both BCoV/Ent and BCoV/Resp infected PBMCS groups, compared to the sham infected cells, reflecting a regulatory response to the infection (Fig. 5E). The TGF.-β expression did not show any changes following BCoV infection (Fig. 5G).

**Fig. 5.**
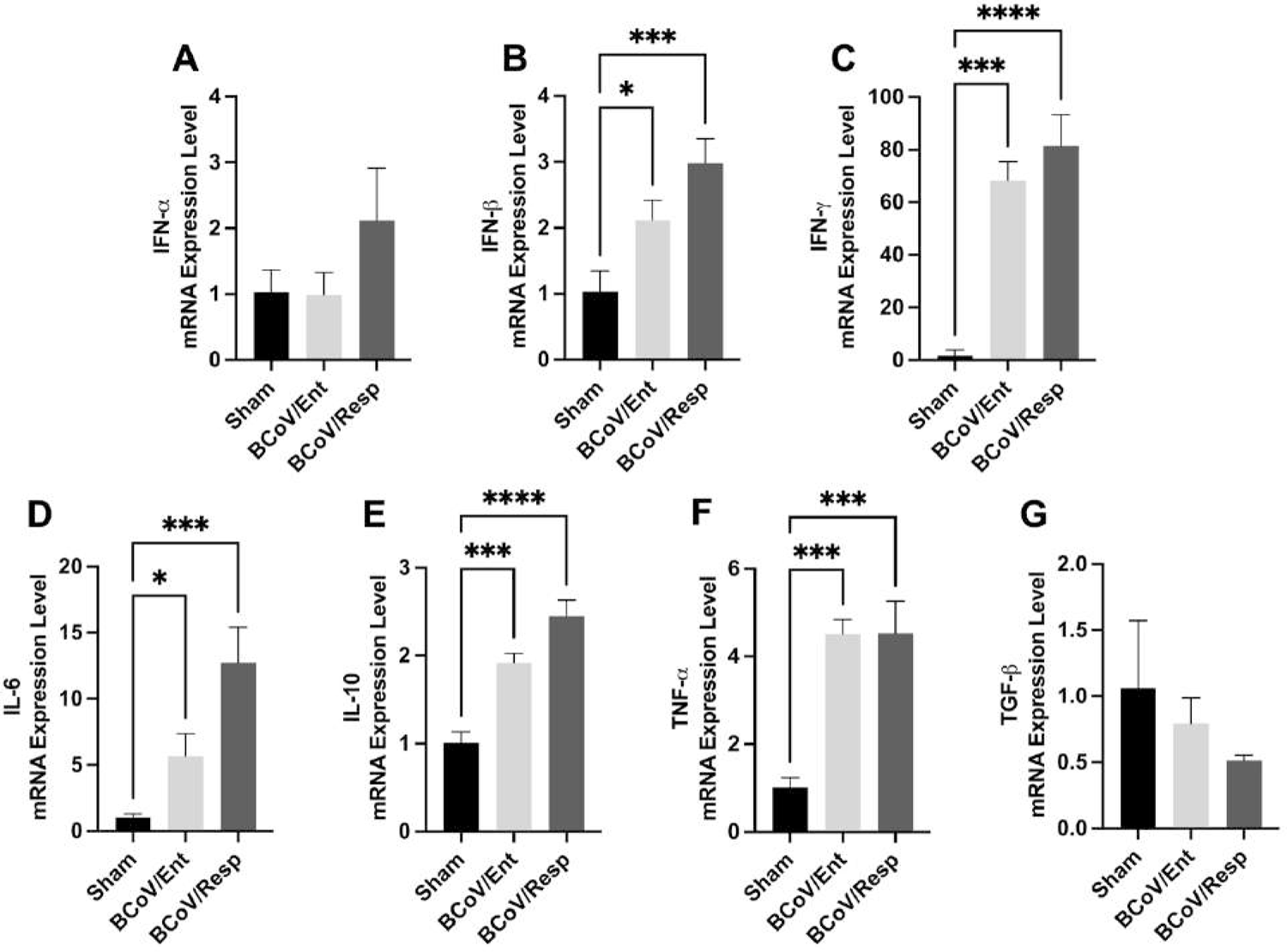
BCoV infection activates the host cell cytokine-like Strome in bovine PBMCs. (A) qRT-PCR analysis was performed to analyze the mRNA expression of Interferon-alpha (IFN-α); (B) Interferon-beta (IFN-β); (C) Interferon-gamma (IFN-γ); (D) Interleukin-6 (IL-6); (E) Interleukin-10 (IL-10); (F) Tumor Necrosis Factor-alpha (TNF-α), and (G) Transformation Growth Factor-beta (T.G.F.-β). Bovine PBMCs were infected with BCoV/Ent and BCoV/Resp at 1 MOI for 72 hours. Total RNAs were isolated from each group to detect the cytokine expression using qRT-PCR. All data were normalized to the sham group.

### 3.6. The BCoV/Ent and the BCoV/Rep infection in the isolated T-cells from the PBMCs induced differential cytokines gene expression levels

Results show that the IFN-α and the IFN-βgene expression levels were upregulated in both the BCoV/Ent and the BCoV/Resp infected T cells, indicating robust innate immune response activation following the BCoV infection (Fig 6A, 6B). The IFN -γ expression level did not show any significant changes in infected and sham infected groups T lymphocytes (Fig. 6C). The pro-inflammatory cytokines IL-6 expression levels were upregulated in both BCoV infected groups, indicating the inflammatory nature of the host immune response (Fig 6D). While the anti-inflammatory cytokine IL-10 expression level showed significant downregulation in both BCoV/Ent and BCoV/Resp groups (Fig. 6E). Together, the upregulation of IL-6 and downregulation of IL-10 indicates a more dominant inflammatory response of the BCoV infection in the T -cells isolated from the PBMCs (Fig. 6D, 6E). The TGF-β expression level was upregulated in the BCoV/Resp group while it was downregulated in the BCoV/Ent group, suggesting different strategies in tissue homeostasis in response to the infection with different BCoV isolates (Fig. 6G).

**Fig. 6.**
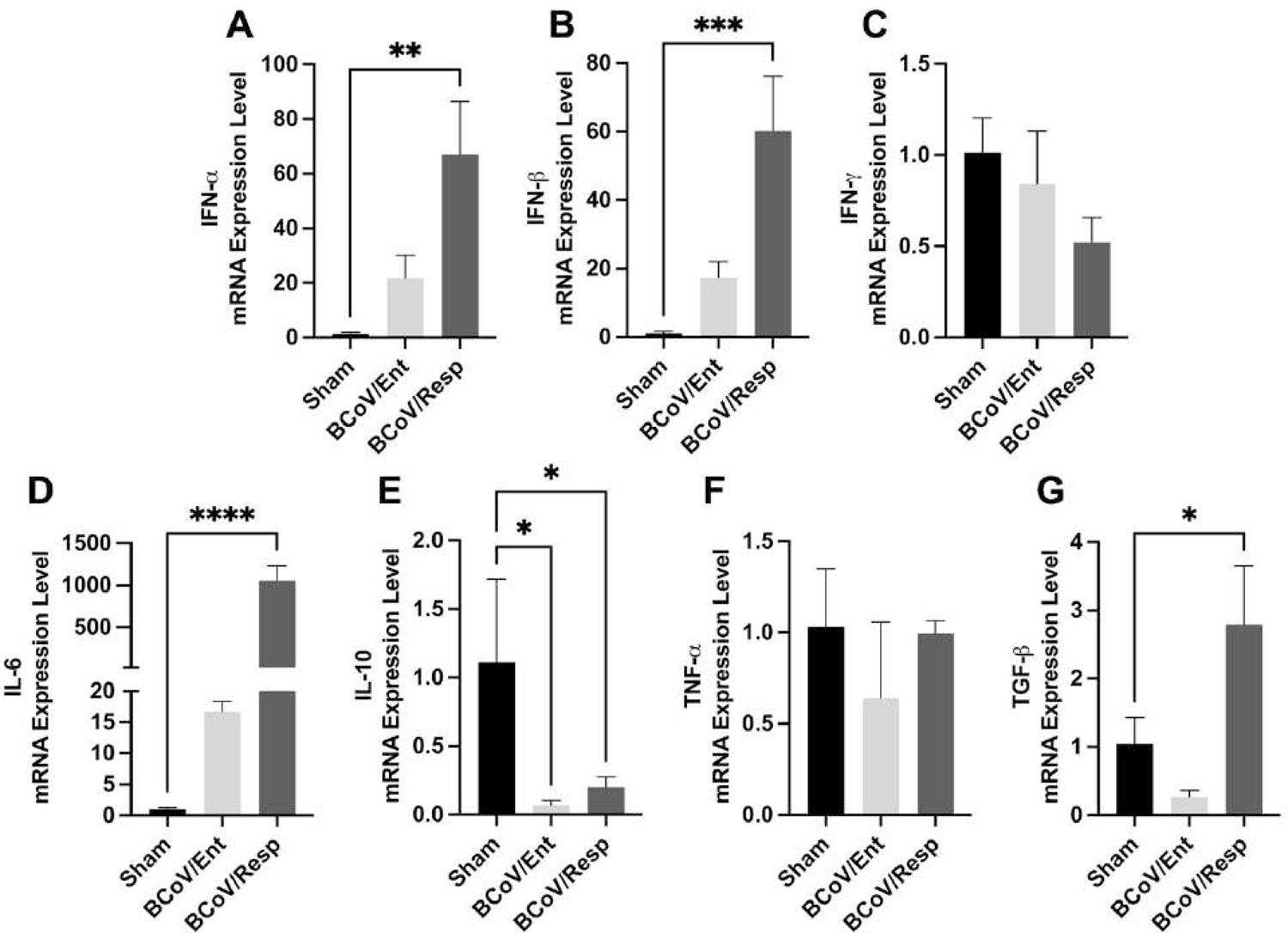
Cytokine gene expression profiles in the T lymphocytes infected with BCoV (A) The qRT-PCR assay was performed to analyze the mRNA expression of IFN-α; (B) IFN-β; (C) IFN-γ; (D) IL-6; (E) IL-10; (F) TNF-α, and (G) T.G.F.-β. Bovine T lymphocytes were isolated from PBMCs and infected with BCoV/Ent and BCoV/Resp at (MOI=1) for 72 hours. The total RNAs were isolated from each group to detect the cytokine expression using qRT-PCR. All data were normalized to the sham group.

### 3.7. Confirmation of the BCoV infection in the bovine splenocytes and their isolated T cells under the *ex vivo* conditions

The bovine splenocytes were isolated from bovine spleen tissue and infected with either the BCoV/Ent or the BCoV/Resp isolates at (MOI=1) for 72 hours. The real-time PCR results demonstrated that the BCoV/Ent and the BCoV/Resp isolates significantly infected the bovine splenocytes under the *ex-vivo* conditions (Fig. 1A). The genome copy numbers in the case of the BCoV/Ent infection demonstrated approximately 1.54 folds higher than that in the case of the BCoV/Resp infected splenocytes (Fig. 7A). The BCoV replication in the isolated bovine splenocytes was also confirmed by the Western blot using the BCoV-S and BCoV-N antibodies (Fig. 7B). Furthermore, to test the ability of BCoV to infect the bovine T lymphocytes isolated from the bovine splenocytes, the qRT-PCR assay assessed the BCoV genome copy numbers. Results confirmed high BCoV genome copy numbers in the case of the BCoV/Ent and BCoV/Rep infected spleen T cells (Fig. 7C). The genomic viral loads in the case of the BCoV/Ent infected group were 61-fold higher than the BCoV/Resp groups, indicating that BCoV/Ent infection is more prominent on splenocytes isolated T lymphocytes (Fig. 7C).

**Fig. 7.**
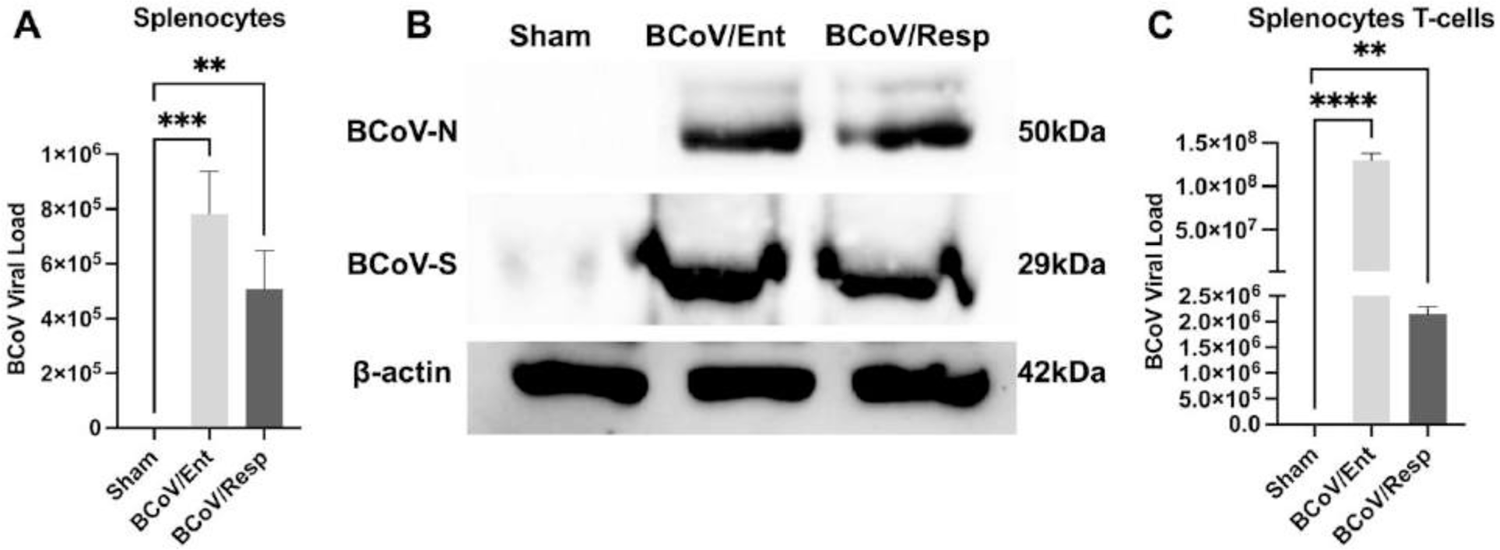
BCoV infects bovine splenocytes and their isolated T-lymphocytes under the *ex-vivo* conditions. (A) The BCoV genome copy number analysis in the bovine splenocytes at 72 hpi through qRT-PCR analysis; (B) Western blot analysis demonstrating BCoV-nucleocapsid (BCoV-N) protein, BCoV-spike (BCoV-S) protein, and β-actin protein expression in splenocytes after 72 hpi. (C) qRT-PCR analysis indicating BCoV genomic viral load in T-cells isolated from splenocytes at 72 hpi.

### 3.8. Confirmation of the BCoV infection/replication in the bovine splenocytes and their isolated T cells by the immunofluorescence under the *ex vivo* conditions

The Immunofluorescence assay confirmed the BCoV infection in bovine splenocytes and their isolated T lymphocytes under *ex-vivo* conditions. The green fluorescent signals, indicative of BCoV Nucleocapsid protein expression, were prominently observed in the BCoV-infected splenocytes (Fig. 8A – 8F). The BCoV-N protein intensities of the green fluorescence were similar in splenocytes infected with either the BCoV/Ent or the BCoV/Resp isolates (Fig. 8E, 8F). Subsequently, the T lymphocytes isolated from splenocytes were examined for the BCoV-N expression levels following BCoV infection under florescent microscopy (Fig. 8G – 8N). Results showed that the CD3 T lymphocytes confirmed the infection with the BCoV, with a similar infection rate observed in both the BCoV/Ent or the BCoV/Resp groups (Fig 8J, 8N). These findings confirm a consistent infectivity pattern of the BCoV/Ent and the BCoV/Resp isolates in the bovine splenocytes.

**Fig. 8.**
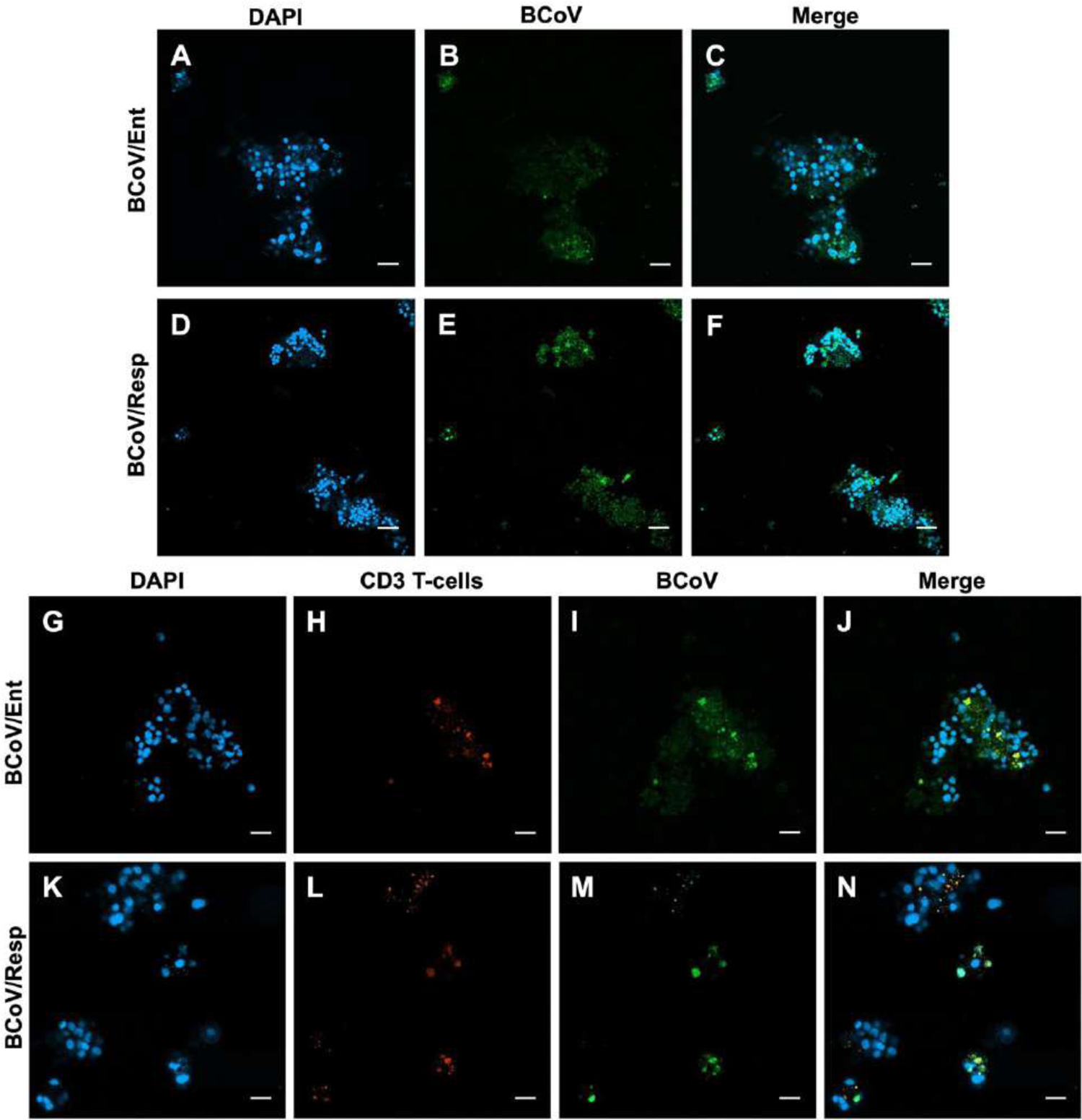
Results of the immunofluorescence assy of the bovine splenocytes and their isolated T lymphocytes after BCoV infection (A-C) The IF staining of splenocytes infected with BCoV/Ent group; (A) DAPI staining of the nuclei in blue, (B) BCoV staining in green color, and (C) Merge showing BCoV binding to the splenocytes. (D-F) IF staining of splenocytes infected with BCoV/Resp group; (D) DAPI staining of the nuclei in blue, (E) BCoV staining in green color, and (F) Merge showing BCoV binding to the splenocytes. (G-J) IF detection of T lymphocytes isolated from splenocytes with BCoV/Ent isolate; (G) DAPI staining the nuclei in blue, (H) CD3 T-cells staining in red color, (I) BCoV staining in green color, (J) and the merged image showing co-localization of T-cells with BCoV. (K-N) IF detection of T lymphocytes isolated from splenocytes with BCoV/Resp isolate; (K) DAPI staining the nuclei in blue, (L) CD3 T-cells staining in red color, (M) BCoV staining in green color, (N) and the merged image showing co-localization of T-cells with BCoV. All the IF images were taken at 20x magnification.

### 3.9. BCoV Infection modulates the proliferation of the CD4/CD8 T-Cells, and the IFN-**γ** production in the bovine splenocytes under the ex vivo conditions

The impacts of the BCoV/Ent or the BCoV/Resp infection on the CD4 and the CD8 T lymphocytes were evaluated in the sham, BCoV/Ent, and BCoV/Resp infected splenocytes via flow cytometry analysis. The histogram representing the fluorescence intensity of the CD4 T-cells demonstrated a notable shift to the left in both the BCoV/Ent and the BCoV/Resp infected groups (Fig. 9A). The MFI of the CD4 T-cells was significantly lower in both infected groups, compared to the sham group (Fig. 9B), suggesting the suppression of the CD4 T-cells in response during the BCoV/Ent and BCoV/Resp infection in the bovine splenocytes. The CD8 T-cell population exhibits a distinct shift to the right in the BCoV/Ent infected group (Fig. 9C). The MFI of the CD8 T-cells was also significantly high in the case of the BCoV/Ent infected group, compared to the sham (Fig. 9D), suggesting the activation of the CD8 T-cells in response to BCoV/Ent infection. The CD4/CD8 ratio showed an inconsistent pattern between the sham and the BCoV-infected groups (Fig. 9G). Specifically, in the case of the BCoV/Ent group, the CD4 T-cell percentage was lower by about 0.88 folds, while the CD8 T-cells were higher by about 1.88 folds than the sham-infected group (Fig. 9G). In the case of the BCoV/Resp group, the levels of the CD4 T-cells were downregulated, but no difference was observed in the CD8 T-cell population compared to the sham group (Fig. 9G). The IFN-γ gene expression in splenocytes showed no significant changes between the sham and the BCoV-infected groups of cells (Fig. 9E, 9F). These results indicate that BCoV/Ent replication/infection was more efficient in splenocytes, suppressing the host CD4 T helper cells and activating the CD8 cytotoxic T-cells.

**Fig. 9.**
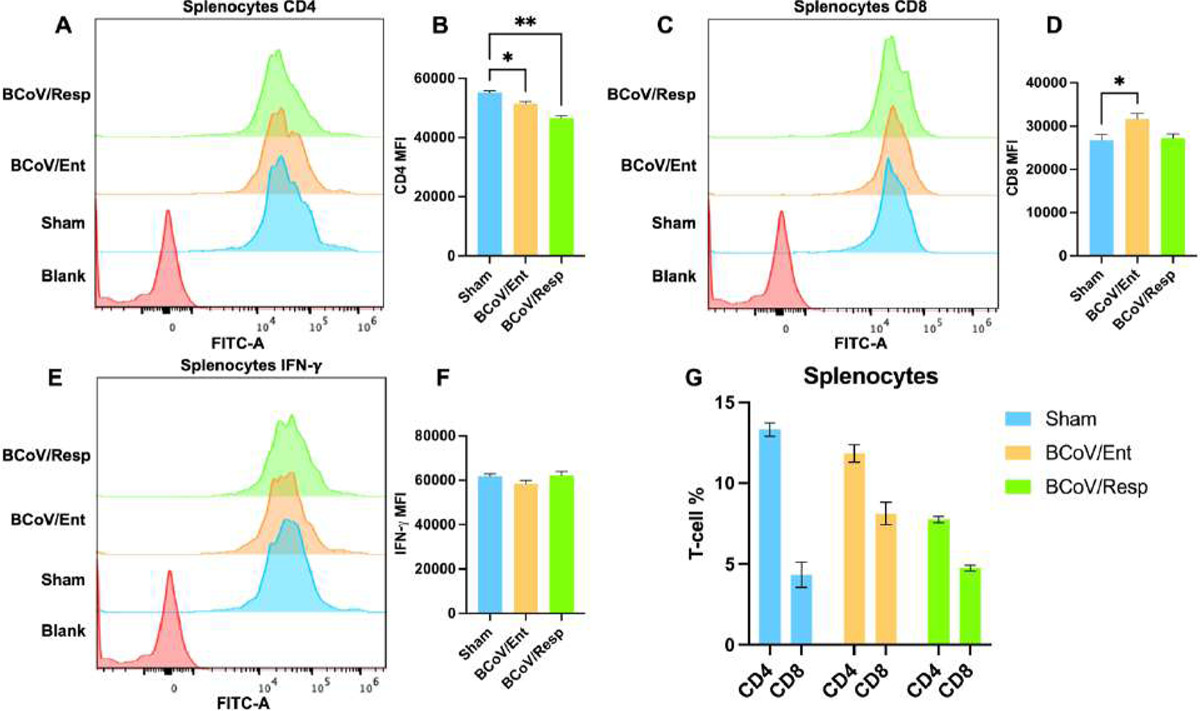
Impacts of the infections on Bovine Splenocyte immunophenotyping. (A) Flow cytometry analysis showing histogram and (B) MFI. of CD4 T-cell population in sham, BCoV/Ent, and BCoV/Resp groups in bovine splenocytes. (C) Flow cytometry analysis showing histogram and (D) MFI. of CD8 T-cell population in sham, BCoV/Ent, and BCoV/Resp groups in bovine splenocytes. (E) Flow cytometry analysis showing histogram and (F) MFI. of IFN-γ in sham, BCoV/Ent, and BCoV/Resp groups in bovine splenocytes. (G) CD4 to CD8 ration in sham, BCoV/Ent, and BCoV/Resp groups in bovine splenocytes.

### 3.10. BCoV Infection differentially regulates the CD4/CD8 Ratios in the T lymphocytes isolated from bovine splenocytes under *ex-vivo* Conditions

The total T lymphocytes were isolated from bovine splenocytes and infected with BCoV/Ent and BCoV/Resp isolates at (MOI=1) for 72 hours and analyzed using flow cytometry. The results demonstrated an obvious shift to the left of the histogram of the CD4 T-cells in both the BCoV/Ent and the BCoV/Resp infected groups, compared to the sham (Fig. 10A). The M.F.I. of the CD4 T-cells was also significantly suppressed in both BCoV -infected groups compared to the sham group (Fig. 10B). The CD8 T-cell population exhibited a distinct shift to the right in both BCoV infected groups of spleen T cells (Fig. 10C). The MFI of the CD8-T-cells were also higher in both the BCoV/Ent and the BCoV/Resp infected groups, compared to the sham group (Fig. 10D), suggesting the upregulation of the CD8 T-cells in response to the BCoV infection. The percentages of the CD4 T-cells showed marked suppression in both BCoV-infected groups (Fig. 10G). At the same time, the CD8 T-cell percentages remain constant in the sham and the BCoV-infected groups (Fig. 10G). The IFN-γ gene expression levels showed no changes in the sham, and the BCoV-infected groups in splenocytes isolated T lymphocytes (Fig. 4E, 4F).

**Fig. 10.**
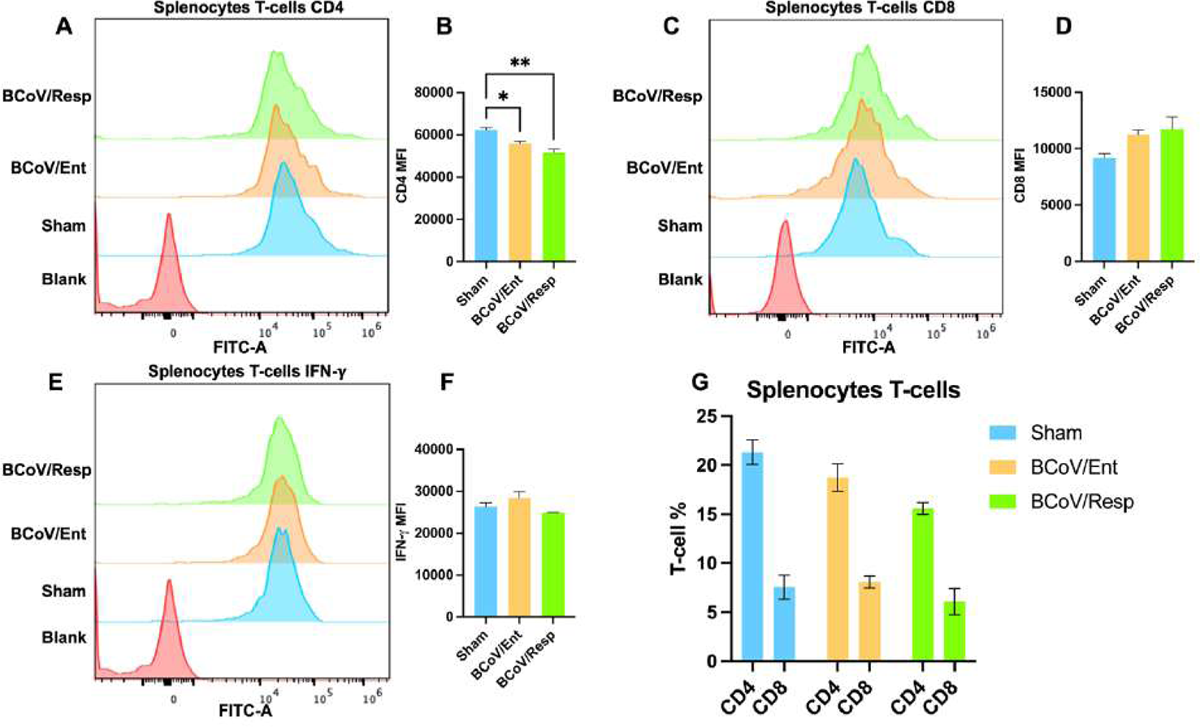
The differential impacts on the immunophenotyping of the BCoV/Ent and the BCoV/Resp isolates infections on the bovine splenocyte and their isolated T lymphocytes. (A)The flow cytometry analysis showing histogram and (B) MFI. of CD4 T-cell population in sham, BCoV/Ent, and BCoV/Resp groups in T Lymphocytes isolated from bovine splenocytes. (C) Flow cytometry analysis showing histogram and (D) MFI. of CD8 T-cell population in sham, BCoV/Ent, and BCoV/Resp groups in T Lymphocytes isolated from bovine splenocytes. (E) Flow cytometry analysis showing histogram and (F) MFI. of IFN-γ in sham, BCoV/Ent and BCoV/Resp groups in T Lymphocytes isolated from bovine splenocytes. (G) CD4 to CD8 ratio in sham, BCoV/Ent, and BCoV/Resp groups in T Lymphocytes isolated from bovine splenocytes.

### 3.11. BCoV infection activates the expression of some host gene cytokines in bovine splenocytes under ex vivo conditions

To examine the bovine cytokines gene expression levels in response to the BCoV infection in bovine splenocytes. The isolated splenocytes were infected with (MOI=1) of either the BCoV/Ent or the BCoV/Resp isolates for 72 hpi. The host cytokine gene expression levels were evaluated using the qRT-PCR assays. Results revealed a significant upregulation of the IFN-β expression levels in the case of the BCoV/Resp infected group, compared to the sham infected groups (Fig. 11B). The IFN-γ expression level was only upregulated in the BCoV/Ent group, compared to the sham (Fig 11C). The pro-inflammatory cytokines IL-6 and anti-inflammatory cytokine IL-10 expression levels did not show any significant changes in sham and the BCoV infected groups (Fig. 11D, 11E). The TNF-α expression level was upregulated in the case of the BCoV/Resp group, compared to the sham (Fig. 11F). The TGF-β expression level did not show any changes in the expression after infection, suggesting a stable regulatory role in immune homeostasis during BCoV infection in bovine splenocytes (Fig. 11G).

**Fig. 11.**
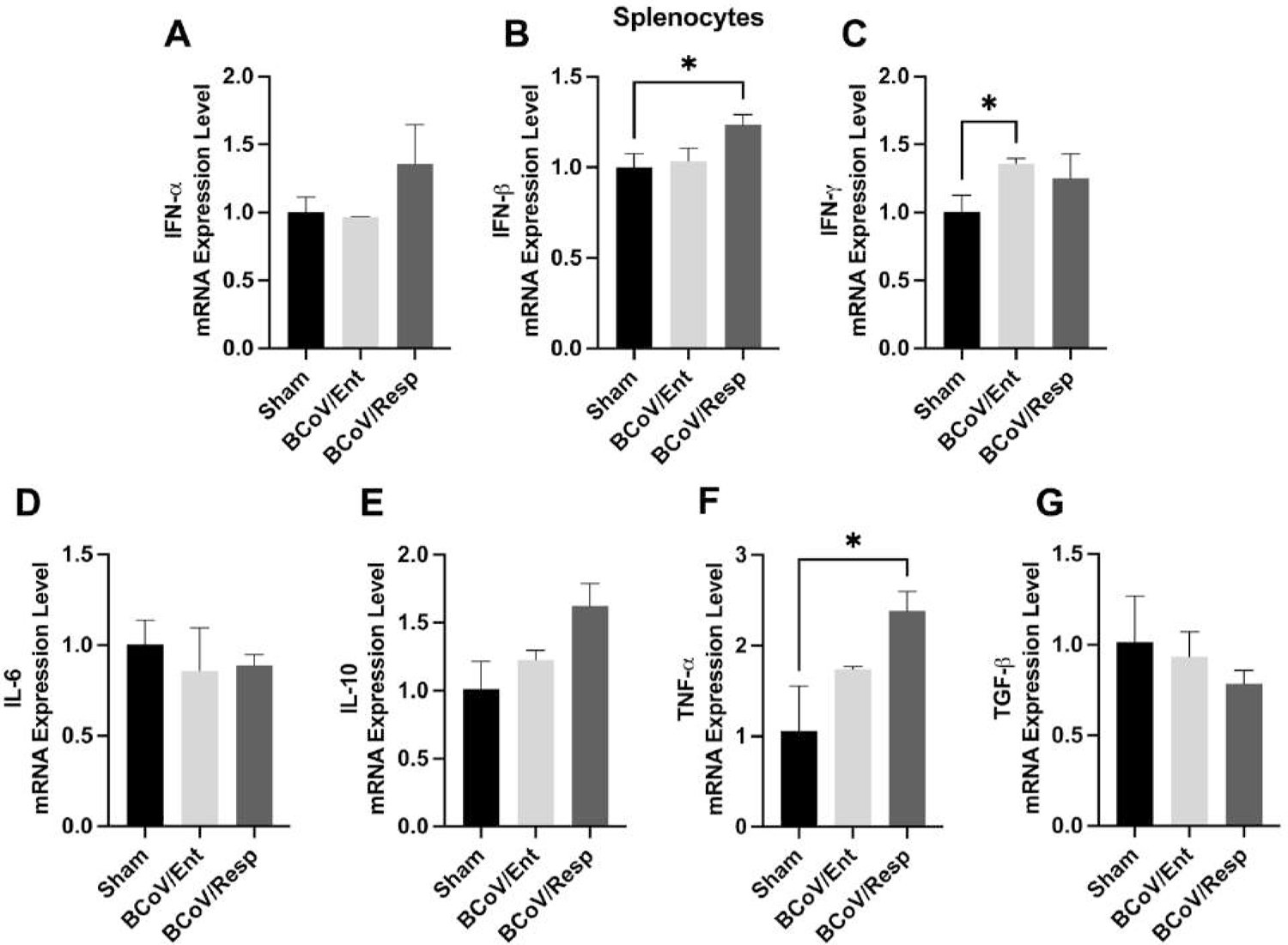
Cytokine gene expression profiles in response to BCoV infection in the bovine splenocytes (A) Results of the qRT-PCR assay to evaluate the mRNA expression of IFN-α; (B) IFN-β; (C) IFN-γ; (D) IL-6; (E) IL-10; (F) TNF-α, and (G) T.G.F.-β in bovine splenocytes. Bovine splenocytes were infected with BCoV/Ent and BCoV/Resp at (MOI=1) for 72 hours. Total RNAs were isolated from each group to detect the cytokine expression using qRT-PCR. All data was normalized to the sham group.

### 3.12. BCoV infection activates some cytokines gene expression in the T lymphocytes isolated from the spleen under ex vivo conditions

To examine the effect of BCoV/Ent and BCoV/Resp isolate on the T lymphocytes isolated from bovine splenocytes, a comprehensive analysis of host cytokines expression was performed under *ex-vivo* conditions. The IFN-α and IFN-β expressions were upregulated in the BCoV/Ent group, indicating the activation of the innate immune responses (Fig 12A, 12B). The IFN-γ expression level did not show any significant changes in the BciV infected and sham groups of T lymphocytes isolated from the spleen (Fig. 12C). The pro-inflammatory cytokines IL-6 expression level was upregulated only in the case of the BCoV/Ent infected group, compared to the sham (Fig 12D). Meanwhile, both the anti-inflammatory cytokine IL-10 and the TNF-α expression levels did not show any significant changes between sham and the BCoV infected groups (Fig. 12E, 12F). The T.G.F.-β expression level was upregulated in the case of the BCoV/Ent infected group, while no significant change in its expression level was observed in the case of the BCoV/Resp infected group (Fig. 12G). The upregulation of the expression levels of the IFN-α, the IFN-β, the IL-6, and the TGF-β in the case of the BCoV/Ent infected group suggests that enteric isolate of BCoV elicited a more robust host immune response compared to BCoV/Resp infected group.

**Fig. 12.**
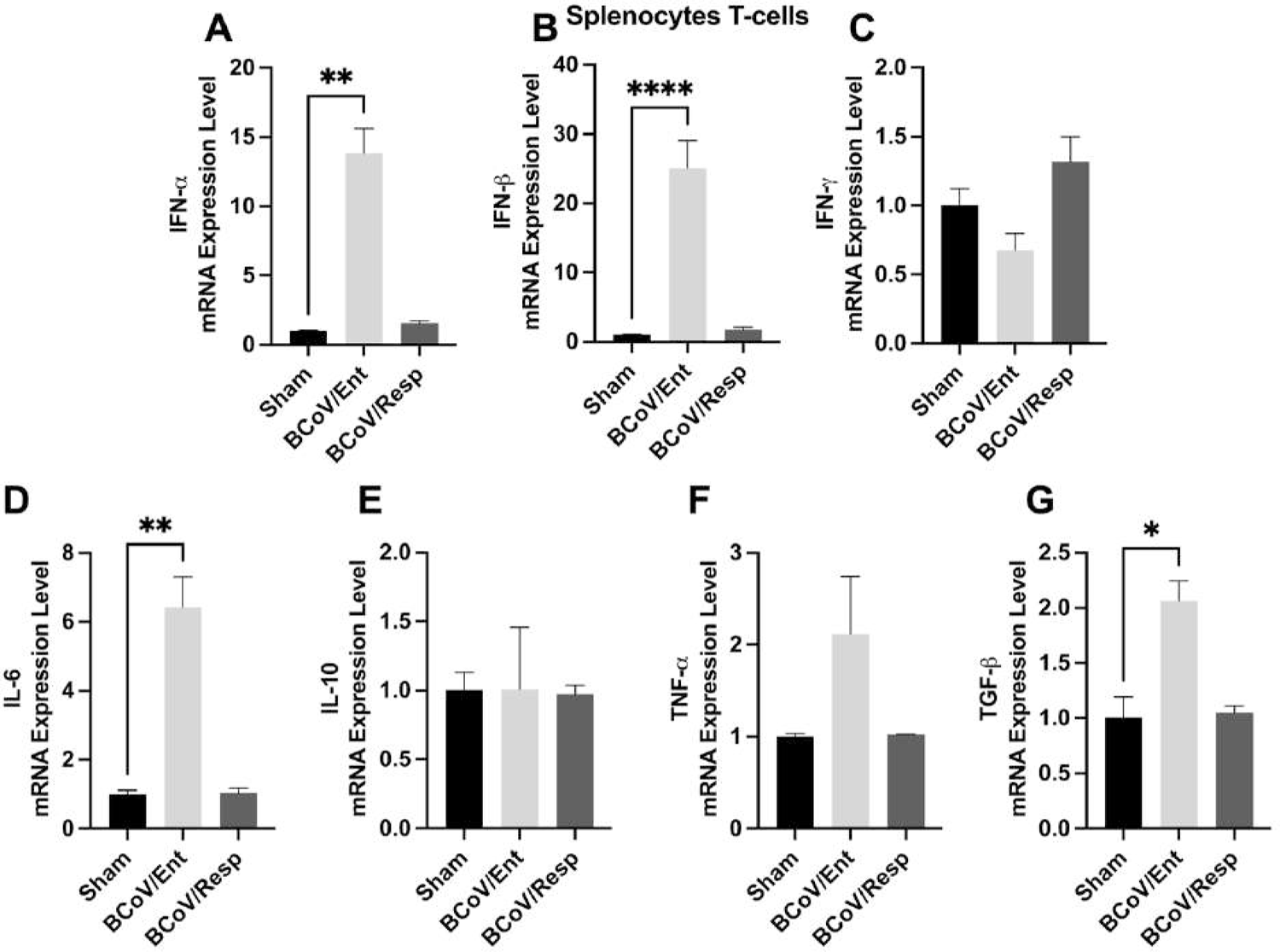
Cytokine gene expression profiles in response to BCoV infection in the T cells isolated from bovine splenocytes (A) Results of the qRT-PCR assay to evaluate the mRNA expression of IFN-α; (B) IFN-β; (C) IFN-γ; (D) IL-6; (E) IL-10; (F) TNF-α, and (G) T.G.F.-β in T lymphocytes. Bovine T lymphocytes were isolated from splenocytes and infected with BCoV/Ent and BCoV/Resp at (MOI=1) 72 hours. The total RNAs were isolated from each group to detect the cytokine expression using qRT-PCR. All data were normalized to the sham group.

## 4. Discussion

Bovine coronavirus (BCoV) infection is ubiquitous, especially in naïve animals during the first three months of age (43). Very little information is available about the humoral immune responses against BCoV, and information about cell-mediated immunity (CMI) during BCoV infection in the infected cell is very scarce. One study was conducted to evaluate the efficacy of the administration of the BCoV live attenuated vaccine in utero to the fetus during the last stage of pregnancy (44). The rationale behind this study was to stimulate the fetus’s immune response to counteract the natural BCoV infection in early postpartum life. The vaccinated animals had a high IgA, IgG, and IgM titer. The vaccinated animals were protected against diarrhea and enteritis compared to the sham-vaccinated animals (44). A recent studies showed that the over-expression of the BCoV-N protein suppresses the IFN-β production by inhibiting the RIG-1 pathway (45). A recent study showed the upregulation of IL-8 chemokine following the challenge of some BCoV seronegative cattle (46).

The peripheral Blood Mononuclear Cells (PBMCs) are isolated from peripheral blood, and they include lymphocytes (T cells, B cells, and NK cells) (47). The spleen is the second lymphoid organ that performs a variety of immunological functions in the body, including red blood cell clearance from pathogens (48). Several studies have been conducted to explore the efficacy of viruses in infecting PBMCs under *ex-vivo* conditions. Messias et al. 2019., showed that PBMCs isolated from healthy donors can be infected by the ZIKA virus under ex-vivo conditions (49). This study showed that PBMCs derived from healthy cattle can be infected with either the BCoV/Ent or the BCoV/Resp isolates under ex-vivo conditions. Furthermore, splenocytes isolated from healthy bovine spleens can also be infected with BCoV/Ent and BCoV/Resp isolates under ex-vivo conditions. Interestingly, our results clearly show that the BCoV replicates more dominantly in the bovine splenocytes (Fig. 7A) compared to the isolated bovine PBMCs (Fig. 1A).

The human PBMCs contain (70 – 90%) lymphocytes, and 70 – 85% of these populations are CD3 T-cells. The CD3 T-cells are complex proteins that carry the T cells’ receptors (TCR). It is expressed in all T cell populations. However, CD4 and CD8 are co/receptors for T cells (50). The CD4/CD8 ratios in cattle ranged from 0.5-2. Monitoring the alteration in the CD4/CD8 ratios is very important to assess the immune status in humans and animals. This CD4/CD8 ratio gives a good indication of the health status of the target host, including cattle (51, 52). The naïve spleen activates memory T-cells in response to cognate antigen and T-dependent B-cell germinal center reactions, producing antibodies upon antigen recognition (53). Therefore, in the present study, T lymphocytes were isolated from the PBMCs and the bovine splenocytes and were infected with either the BCoV/Ent and BCoV/Resp isolates. The BCoV infection rate was notable in the isolated T lymphocytes from the PBMCs and the bovine splenocytes (Fig. 1C, 7C).

The SARS-CoV-2 infects T lymphocytes, especially CD4 T-cells, causing pronounced T-cell death and potentially contributing to lymphopenia in the COVID-19 patients (54). Another study demonstrated that cross-protection against SARS-CoV-2 elicited by HCoV-OC43 depends on the CD4 T-cell through the T-cell mediated mechanism (55). In this study, we observed the CD4 T-cell population was upregulated in PBMCs and T lymphocytes after the BCoV/Ent infection, while the CD8 T-cell population was downregulated after infection in both groups (Fig. 3, 4). This suggests that BCoV/Ent infection leads to a more severe response in the PBMCs and their isolated T lymphocytes, activating the host CD4 T helper cells while inhibiting the CD8 cytotoxic T-cell responses. Conversely, in splenocytes and their isolated T lymphocytes, the CD4 T-cell population was downregulated, and the CD8 T-cell population was upregulated following BCoV infection (Fig. 9, 10). This indicates the activation of the cytotoxic CD8 T-cells in splenocytes in response to the BCoV infection, demonstrating the ability of splenocytes to mount a response and potentially inhibit BCoV infection through a CTL-mediated mechanism. Collectively, these results suggest that the immune response to BCoV infection involves tissue-specific modulation of T-cells. The upregulation of CD4 T-cells in PBMCs may indicate an early immune activation, while the upregulation of CD8 T-cells in splenocytes indicates a targeted cytotoxic response to BCoV infection. Several studies highlighted the crucial role of the bovine T lymphocytes in response to various viral infections (6, 56). The related studies demonstrated the presence of the CD8 and the CD4 memory T-cells and their associated cytokines in the BVDV seropositive cattle (23). Similarly, another study showed the cytotoxic T lymphocytes (CTL) in the PBMCs from immunized cattle were capable of killing the autologous BVDV-infected antigen-presenting cells upon in-vitro re-stimulation (57). In cattle’s, the roles of the major T-cells subsets, such as T killer (CD8 T-cells), T helper (CD4 T-cells, including subsets Th1, Th2, and Th17), and T regulatory cells (CD4+/CD8+CD25+) have been investigated in various infections and post-vaccine immune responses (58, 59). The Th1 cells responding to cattle parasites (bovine tuberculosis) can produce IL-2 and IL-10, in addition to IL-4 and IFN-γ production (60). While the Th1 cells only produce IFN-γ and IL-2 in response to viral infection (bovine leukemia virus) (14). In contrast, Vanden Bush demonstrated that Mycoplasma bovis (Bacterial infection) did not result in IFN-γ production in the tested PBMCs (61). In this study, the expression levels of the IFN-γ and the IL-10 were upregulated in the PBMCs infected with BCoV (Fig. 5C, 5E), while the expression levels of both the IFN-γ and the IL-10 were downregulated in the T lymphocytes isolated from PBMCs (Fig. 6C, 6E). BCoV infection inhibited the T helper cell regulation in the PBMCs, while no significant changes were observed in the isolated bovine splenocytes infected with the BCoV. The TGF-β, another crucial anti-inflammatory cytokine, was only upregulated in the T lymphocytes isolated from the PBMCs or the splenocytes infected with BCoV (Fig, 6G, 12G).

The consistent upregulation of the expression levels of the IFN-α and the IFN-β in the T lymphocytes isolated from the PBMCs and splenocytes highlights the activation of the innate immune response during BCoV infection in the *ex vivo* models (Fig. 6A, 6B, 12A, 12B). The expression levels of the pro-inflammatory cytokine IL-6 were also upregulated in both the BCoV/Ent and the BCoV/Resp infected groups in both the PBMCs and their isolated T lymphocytes (Fig. 5D, 6D). In contrast, IL-6 expression level was only upregulated in the case of the BCoV/Ent infected group in the splenocytes and their isolated T lymphocytes (Fig. 11D, 12D). It was reported that SARS-CoV-2 infection also triggers a robust IL-6 production in the host, which results in cytokine Strome production (62). These results indicate the importance of type-1 interferon in the early innate immune response in bovine PBMCs and splenocytes during the BCoV infection. Furthermore, IL-6 expression is more dominant in PBMCs compared to the bovine splenocyte.

## Conclusions

This study confirms for the first time the successful infection and replication of the BCoV enteric and respiratory isolates in the PBMCs, their isolated T Cells, the bovine spleen, and their isolated T cells. The high viral genome copy numbers supported the successful BCoV replication in the PBMCs and the splenocytes, the BCoV proteins (S and N) expressions, and the demonstration of the BCoV-N protein in the infected cells by the IFA. We also confirm a consistent pattern of type 1 interferon (IFN-α and IFN-β) production during BCoV infection in the ex vivo models using the PBMCS and the splenocytes, indicating a significant innate immune response activation in both cell types. Furthermore, the pro-inflammatory cytokine IL-6 and the anti-inflammatory cytokine IL-10 expression levels exhibited distinct expression patterns, reflecting complex regulatory dynamics in response to the BCoV infection. Furthermore, the differential modulation of CD4 and CD8 T-cells indicates the early activation of CD4 T helper cells in bovine PBMCs in response to BCoV infection. These findings enriched our understanding of the host immune response during BCoV infection. It also provides critical insights for developing novel immunotherapies and vaccines against BCoV infection in cattle, improving cattle health and disease management.

## Acknowledgments

We thank Dr. Aspen Workman from the Animal Health Genomics Research Unit, USDA, ARS., U.S. Meat Animal Research, for kindly providing the bovine coronavirus respiratory isolate used in this study. We also thank Drs. James Q. Robinson and Sukolrat Boonyayatra, for their technical and logistical support with the sample tissue collection.

## Data availability statement

All the data are available upon request from the corresponding author.

